# Deciphering Photosynthetic Protein Networks: A Crosslinking-MS Strategy for Studying Functional Thylakoid Membranes

**DOI:** 10.1101/2025.10.07.681025

**Authors:** Nicolas Frances, Cecile Giustini, Giovanni Finazzi, Myriam Ferro, Pascal Albanese

## Abstract

Photosynthesis, which sustains life on Earth, depends on organized and yet adaptable protein assemblies embedded in specialized membranes known as thylakoids. Understanding how these complexes interact and reorganize within functional photosynthetic membranes is essential to reveal the molecular basis of energy conversion in cells. Here, we present an improved crosslinking mass spectrometry strategy that captures native protein interactions in photosynthetically active thylakoid membranes from *Arabidopsis thaliana* and *Spinacia oleracea*. By monitoring photosynthetic performance during crosslinking, we show that electron transport remains active, allowing structural analysis under physiological conditions. Moreover, we show that trimethylphenylammonium chloride (TMPAC) as an adjuvant charged compound does not impair physiological activity, while boosting and diversifying crosslink identifications. Mapping crosslinks onto known structures confirms the integrity of major photosynthetic complexes and uncovers previously uncharacterized assemblies involving regulatory and structural proteins. Integration with structural modeling and interaction network analysis identifies novel protein players within the photosynthetic machinery, providing molecular insights into their potential roles. This approach offers a broadly applicable framework for studying membrane protein organization and dynamics in functional bioenergetic systems.

**Significance Statement:** Photosynthesis relies on dynamic protein interactions within thylakoid membranes, yet capturing these networks under physiological conditions remains challenging. We establish a crosslinking mass spectrometry workflow that maps native protein interactions in plant thylakoids while preserving photosynthetic activity. By bridging structural and functional biology, this approach enables *in situ* exploration of membrane protein networks and advances our understanding of photosynthetic regulation

## Introduction

Throughout billions of years of evolution, living cells have developed a wide range of specialized subcellular compartments. Within each, proteins perform biochemical functions as part of macromolecular assemblies, these in turn form a large network of molecular interactions within a cell, referred to as its interactome. Energy metabolism in eukaryotic cells also relies on a dynamic network of protein complexes embedded in the membranes of specialized organelles: the ubiquitous mitochondria and, in oxygenic eukaryotic phototrophs, the chloroplasts. Oxygenic photosynthesis has driven nearly all life forms on Earth since over 3.5 billion years, using sunlight as the energy source, thus coping with its variability in intensity and quality across nearly every habitat (Blankenship 2010). This remarkable adaptability lies with complex regulatory networks inside the chloroplast thylakoid membranes where highly conserved catalytic cores, the photosystems (PS) I and II, work in series with the Cytochrome b_6_f and ATP synthase complexes (Eberhard et al. 2008). Opposed to the overall evolutionary conservation of these major complexes, several proteins interacting with them, stably or transiently, have been the evolutionary key in photosynthetic adaptation, both for harvesting sunlight in form of external antenna systems (Croce and van Amerongen 2020) and mitigating its harmful effects through photoprotective mechanisms (Bassi and Dall’Osto 2021). The cellular interactome can adapt much faster to environmental cues than the transcriptome or the proteome (Bludau and Aebersold 2020), and its changes represent therefore a major target to study how organisms adapt to environmental cues. These organisms power our society: from oil and coal to our main food crops; our energy supply is based on the productivity of photosynthetic life. Consequently, understanding the photosynthetic thylakoid’s interactome dynamics in a near-native and functional state is of primary importance to meet future challenges (Ort et al. 2015).

Multiple techniques developed in recent years have enabled the detection and visualization of protein networks and their architectures, often retaining the cell’s native or near-native environment (McCafferty et al. 2024). These include confocal expansion microscopy, cryo-electron microscopy and tomography (cryo-EM/ET), and mass spectrometry (MS) (Klykov et al. 2022; Kelley et al. 2024). For cryo-EM/ET approaches the major limitations is the need of biochemical purification (Cryo-EM single particle) and sample snap-freezing prior to analysis, sometimes with long procedures and difficult access to state-of-the-art technical facilities (Beck and Baumeister 2016). For MS-based analysis, it’s necessary to denature and digest samples prior to data acquisition (McCafferty et al. 2024). For any approach, limitations are exacerbated when analyzing membrane proteins. It appears clear that a multifaceted approach combining spatial organization with functional and molecular identity will be key to understand biological complexity across scales.

A promising approach to study protein-protein interactions across several scales of complexity is crosslinking (XL) coupled to MS (Chavez et al. 2018; O’Reilly and Rappsilber 2018). The approach relies on the use of chemical crosslinking reagents forming covalent bonds between amino acids in close spatial proximity. Once the two crosslinked peptides are sequenced by MS, they yield a distance restraint, typically between 6 and 35 Å, at the residue level (Belsom and Rappsilber 2021). These are particularly suitable to be fruitfully integrated with cryo-ET and cryo-EM imaging data (Kastritis et al. 2017; O’Reilly et al. 2020; Mosalaganti et al. 2022; Albanese et al. 2020).

The use of XL is not new in plant biochemistry (Baird and Hammes 1976; Henriques and Park 1978), but the increasing sensitivity of MS instruments has improved the yield of reliable identifications. The gain in popularity of XL-MS is indeed mainly due to its relatively ease of use, robustness, increasing availability of crosslinking reagents with adaptable chemistry (Steigenberger et al. 2020). Recent integration with cryo-ET/EM and AI-driven structure prediction tools has further boosted its appeal, as XL-MS can both validate (Burke et al. 2023) and drive these predictions. So far, limiting factors for proteome-wide studies have been mostly the high dynamic range of cellular proteomes (Chavez et al. 2019; Liu et al. 2018), requiring extensive fractionation and long MS acquisition times. In addition, a non-negligible limitation of this approach is that it may not yield physiologically relevant structural information due to the covalent bonds formed during the reaction. Several attempts have been undertaken to identify the sub-cellular interactome of mitochondria (Hevler et al. 2023; Liu et al. 2018) and eukaryotic cells (Kastritis et al. 2017), while only few examples exist for plants chloroplasts/thylakoids (Albanese et al. 2020; Ditz et al. 2024; Weisz et al. 2017) or microalgae. XL-MS provides a powerful approach that can be applied holistically without a predefined target, with incubation times as short as 20 minutes. From a structural modelling standpoint, XL-MS also allows retrieval of defined spatial constraints (in the range between 6-35 Å) to model protein-protein interaction interfaces and dynamics (Mendes et al. 2019; Bar Barroeta et al. 2024). Finally, no XL-MS study has been coupled to functional measurements to prove it can yield physiologically relevant results.

In this study, we developed a fast and reproducible pipeline that improves cross-linking efficiency in functional thylakoid membranes extracted from two plant species: *Spinacia oleracea* (spinach) and *Arabidopsis thaliana.* The main advantage of using functional thylakoid membranes is that we were able to measure the physiological impact of chemical crosslinking *in situ* during the chemical reaction. Although isolated thylakoids do not encompass the complete chloroplast interactome—lacking stromal, envelope, and cytosolic components—they remain a well-established system for investigating photosynthetic membrane organization and electron transport under controlled conditions. Isolated thylakoid membranes revealed fundamental mechanistic insights for several decades (see for instance Andersson and Anderson 1980; Bassi et al. 1985). More recently, expansion microscopy have shown that purified thylakoids largely preserve native membrane architecture (Bos et al. 2023; Berentsen et al. 2025), even though the arrangement of photosynthetic complexes in exploded chloroplasts may not reflect entirely their native arrangement (Wietrzynski et al. 2025). Our goal here is therefore to capture near-native functional states of the thylakoid membrane to identify novel protein interactions in an unbiased manner.

Thylakoid membrane surfaces are densely packed of proteins and partially shielded by surface charges (Kirchhoff 2019). This can impair the accessibility and create electrostatic repulsion for certain crosslinkers (Belsom and Rappsilber 2021). Here, we used an enrichable cross-linking reagent (disuccinimidyl phenyl phosphonic acid, or PhoX - Steigenberger et al. 2019) to generate a robust and reproducible output with hundreds of unique crosslinks while keeping MS-acquisition times bearable and sample processing times within few days for several replicates and conditions. One downside is that PhoX carries a negatively charged phosphonic group, which may reduce its proximity to negatively charged membranes and protein surfaces. To mitigate this physico-chemical constraint, we tested an adjuvant compound: trimethylphenylammonium chloride (TMPAC). As an amphiphilic cation, TMPAC may likely reduce electrostatic repulsion and improve local crosslinker accessibility without acting as a detergent or disrupting membrane integrity.

With the guiding principle of establishing a reliable and reproducible protocol we have tested two commonly used plants, spinach (*Spinacia oleracea*) and *Arabidopsis thaliana* grown in varying light conditions, crosslinked at different concentrations of both crosslinker and total protein. In addition, out of dozens of new protein-protein interactions detected, we provide few examples of possible usage of XL-MS data to model novel functional complexes. We believe that this workflow, combined with *in silico* and predictive AI-based structural modelling, will pave the way for integrative studies on functional membrane systems and bioenergetics organelles, beyond photosynthesis research.

## Results and Discussion

### Establishing a Physiological Benchmark for *In Situ* Structural Proteomics of Functionally Active Thylakoid Membranes

Studying protein interactions within native biological membranes presents a formidable challenge due to the need to preserve the delicate structural and functional context of the system. Multiple crosslinking reagents have been developed to cover specific constraints in interaction distances and aid during MS identification (Belsom and Rappsilber 2021). PhoX is a functionalized crosslinker with a phosphonic group, which allows peptide enrichment by immobilized metal-affinity chromatography (IMAC, Steigenberger et al. 2019). This same negatively charged functional group is however believed to cause its cell-impermeability (Steigenberger et al. 2019), which has been reported to be repelled by cellular membranes (Vedadghavami et al., 2020). This might reduce PhoX crosslinking efficiency due to coulombic repulsion from exposed acidic amino acids (Lyu et Lazár, 2022; Kaňa et Govindjee, 2016). To optimize PhoX application on functional thylakoids, we propose a pipeline incorporating TMPAC, an amphiphilic ammonium salt. Its cationic group may neutralize surface charges, allowing PhoX proximity, and improving protein interaction capture.

To assess whether our crosslinking protocol preserves the physiological integrity, we measured photosynthetic activity proxies: the maximum quantum yield of PSII (Fv/Fm) and Electron Transport Rate (ETR) in thylakoid membranes treated with crosslinkers, with and without adjuvant chemical molecules (*i.e.* TMPAC). We first combined confocal imaging of isolated thylakoids upon chemical crosslinking treatment (with and without TMPAC) with functional measurement of photosynthetic electron flow by chlorophyll fluorescence. This allowed us to establish an optimized protocol for the isolation of functional thylakoids from both model organisms. Purified thylakoids retain intact grana and thylakoid organization and most also retain the envelope (Figure S1), as also shown by recent reports (Bos et al. 2023; Berentsen et al. 2025).

To examine the photosynthetic electron transport upon illumination, we measured the PSII quantum yield while increasing the intensity of continuous light from 8 to 550 μmol photons m⁻² s⁻¹ with steps of 5 minutes each for a total of 40 minutes (Figure 1A, Figure S2). First, a titration of increasing Phox and TMAPC concentrations on both species showed that even very low amounts of crosslinker impair the photosynthetic electron transport (Figure S2). On the other hand, TMPAC concentrations up to 100 mM did not affect the ETR, and showed no significant addictive effect when used in combination with Phox. In both spinach and *Arabidopsis* thylakoids, PhoX at 1 or 2 mM reduced ETR by half, indicating partial inhibition of electron flow (Figure 1A and Figure S2). This effect is likely due to covalent spatial crosslinking of electron transport-associated proteins, and possibly the chemical modification *via* the NHS ester labelling of key lysine residues of PsaC (Fischer 1998) and cytochrome b_6_f (Gong et al. 2000). *Arabidopsis* thylakoids treated with PhoX and TMPAC showed a 52% ± 13% decrease in ETR, only marginally lower than with PhoX alone (44% ± 10%). This reduction occurred without drastic alterations in the PSII quantum yield in the dark (Fig. 1B, and Figure S2), suggesting that the interference in electron flow occurs during illumination, while in the dark the thylakoid membrane organization and PSII structural configuration remain largely preserved. This can be expected since we partially introduce covalent bonds between proteins and electron carriers that need dynamic delocalization for their functioning. We therefore decided to perform the subsequent experiments in dark-adapted conditions.

**Figure 1.**
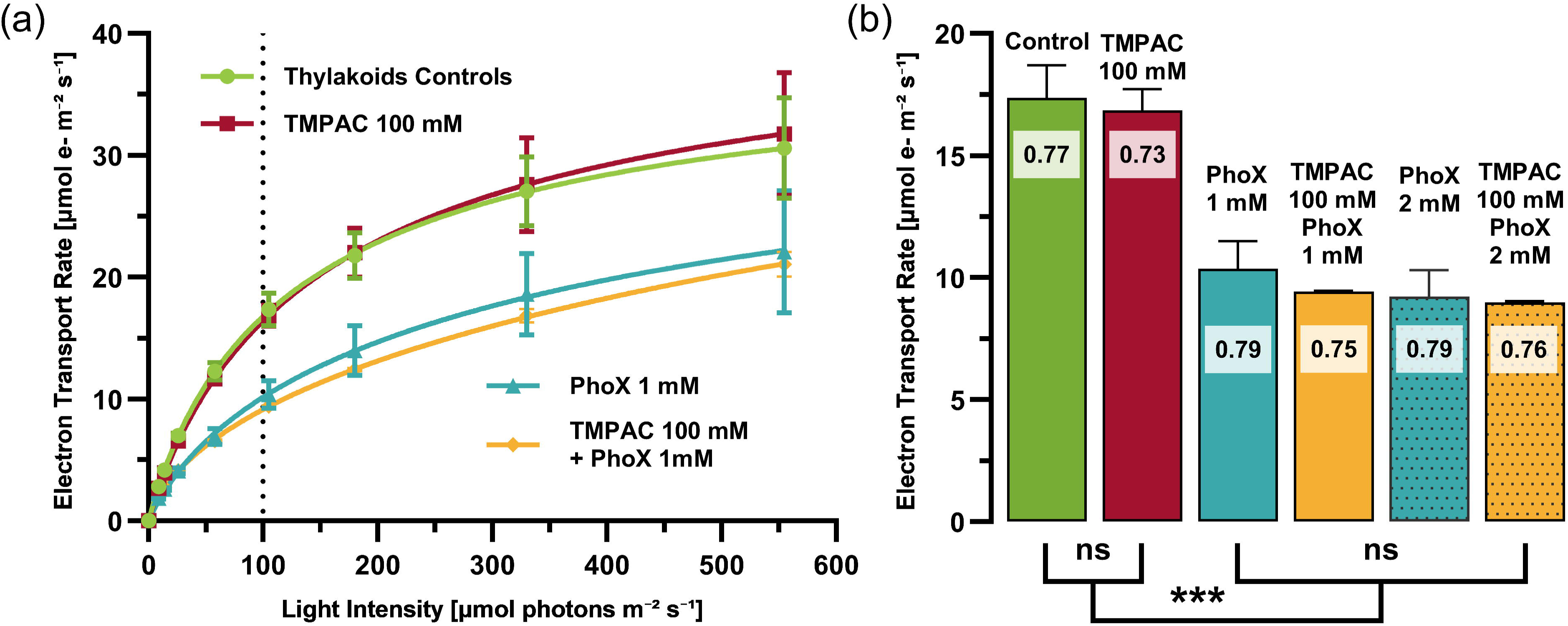
Photosynthetic activity is partially retained during crosslinking of functional thylakoid membranes. (A) Electron transport rate (ETR) curves of spinach thylakoids under increasing light intensity show how crosslinking with PhoX for 40 minutes during the measurement affects photosynthetic performance. (B) ETR values under moderate light (105 μmol photons m⁻² s⁻¹, dashed line in panel A) are plotted for multiple treatment conditions. PSII quantum efficiency in the dark (Fv/Fm) is also reported overlayed to the bar in a textbox (complete values in Figure S2).

Altogether, these physiological measurements confirm the feasibility of the crosslinking approach on functionally active membranes, an essential step forward for membrane systems biology. They demonstrate that crosslinking can be performed on isolated thylakoids in near-native, physiologically active states. The next step is to evaluate the MS readout and the structural data that can be extracted, to detect structurally informative crosslinks under physiologically relevant conditions.

### TMPAC treatment increases the yield of Crosslinked Peptides without introducing a detectable bias

Mass spectrometry analysis of the crosslinked peptides identified a total of 832 and 828 unique crosslinks in spinach (*So*) and *Arabidopsis* (*At*) thylakoids in a total of 8 independent experiments with 3 replicates each (Figure 2a, Table S2). The total number of reproducible crosslinks (XL), identified by crosslinks-spectra matches (CSM) varied between plant species and experimental conditions. Ranging roughly between 200 and 400 reproducible XL, *i.e.* identified in at least 2 replicates (Figure 2a). The number of unique protein-protein pairs in this analysis was 233 and 410 for spinach (So) and *Arabidopsis* (At), respectively (Table S2). Notably, the addition of TMPAC increased the identification of crosslinked peptides across all experimental conditions (Figure 2b). The enhancement ranged a negligible 3.8% up to 55.7% for individual samples, with an average increase of 23.6% across two out of three replicates and 30.6% across three out of three replicates (Figure 2c).

**Figure 2.**
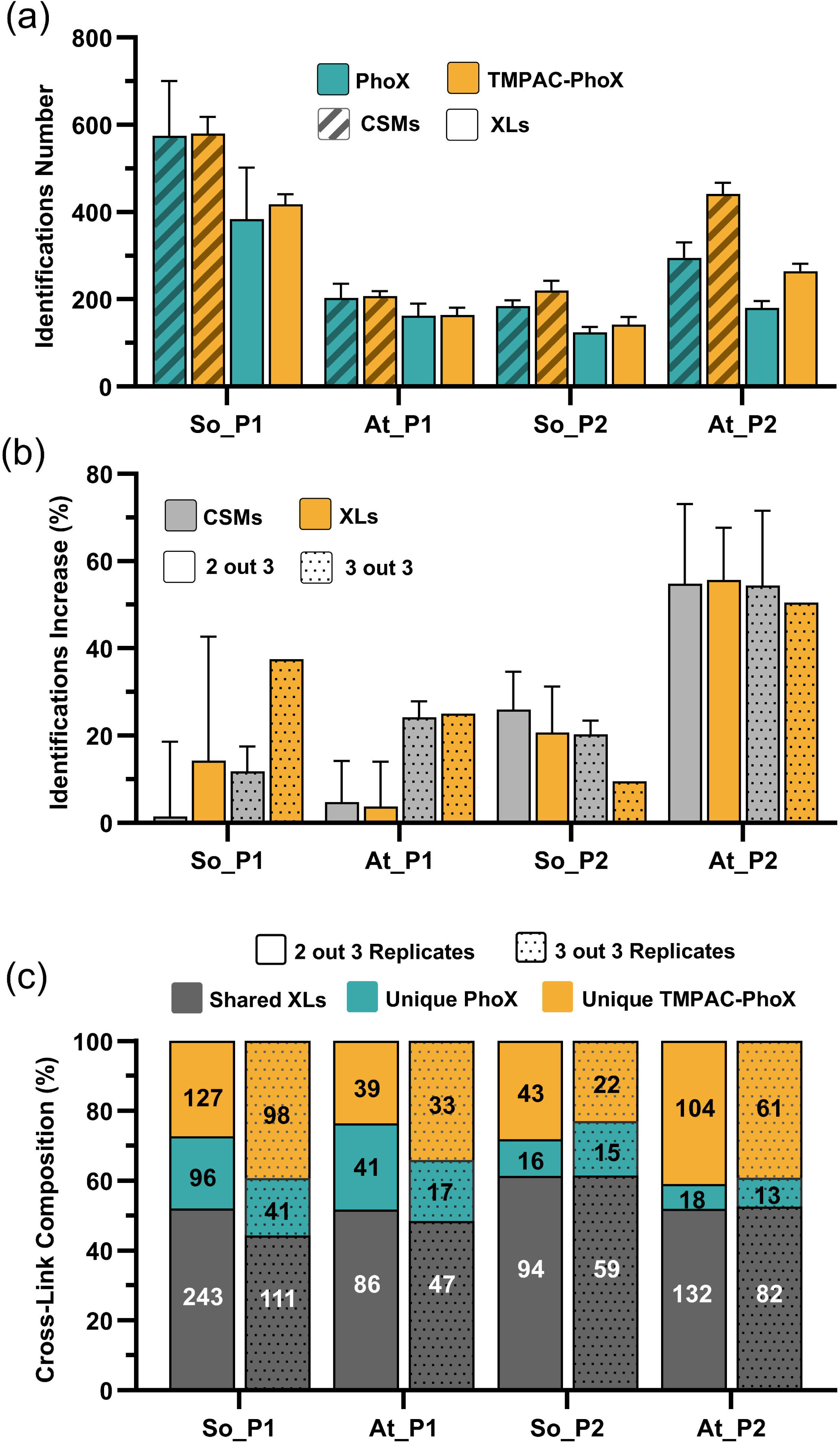
Enhanced detection of protein interactions within functional thylakoid membranes. (a) Crosslink spectrum matches (CSMs; hatched bars) and unique crosslinks (XLs; solid bars) identified from three independent replicates of thylakoid samples from *Arabidopsis thaliana* (At) and *Spinacia oleracea* (So) crosslinked with PhoX for 30 minutes in the presence of TMPAC (orange) or with PhoX alone (teal). Two PhoX concentrations are shown (P1 = 1 mM; P2 = 2 mM). Bars represent mean ± SD (n = 3). (b) Increase in crosslink identifications obtained with TMPAC-PhoX compared to PhoX alone (CSMs in gray and XLs in yellow). Plain bars indicate crosslinks detected in at least 2 of 3 replicates, and dotted bars indicate crosslinks detected in all 3 replicates. Values represent mean ± SD (n = 3). (c) Composition of XLs for each condition showing shared crosslinks between PhoX and TMPAC-PhoX (dark gray), crosslinks unique to PhoX (teal), and crosslinks unique to TMPAC-PhoX (orange). Solid bars indicate crosslinks found in at least 2 of 3 replicates, and dotted bars indicate crosslinks found in all 3 replicates. Source data are reported in Table S1b. Numbers inside bars denote XL counts for each category.

While approximately half of the detected crosslinks are shared between the two treatments (Figure 2c) about one third of the crosslinks could only be identified upon TMPAC treatment when considering highly reproducible crosslinks. Indeed, considering only crosslinks detected in all 3 replicates, the proportion of total XLs and CSMs increased significantly in the TMPAC-treated samples, along with a rise in the number of crosslinks uniquely detected in the presence of TMPAC (Figure 2c). This suggests that TMPAC not only increases crosslinking yield but also enhances its reproducibility.

The treatment however also determines a loss of 10-20% of the identifications uniquely detected when Phox is used alone (FIgure 2c). To further investigate whether this loss is due to the influence of the local sequence environment of TMPAC or PhoX-mediated crosslinking, we analyzed the amino acid composition within a ±3 residue window surrounding each crosslinked lysine (or N-terminal amine) (Figure S3 and Table S3). Across all conditions, hydrophobic residues were the most abundant class, followed by polar and special-case residues, indicating a generally stable sequence context around crosslinked sites. Positively charged residues consistently outnumbered negatively charged residues near the crosslinking position. We could not observe systematic shifts in amino acid class distribution in peptides uniquely crosslinked in the presence of TMPAC (Figure S3c,f,i,l). Furthermore, analysis of crosslink distribution at the protein level did not reveal consistent enrichment toward specific proteins or functional classes (Table S4). Together, these results indicate that TMPAC diversifies crosslink detection without introducing detectable sequence-or protein-level bias.

Overall, we identified hundreds of reproducible crosslinked peptides in a physiologically active sample, and in a rapid manner using the enrichable crosslinker Phox. The addition of TMPAC as an enhancer allowed the detection of 20-30% more crosslinks identified in all replicates, and the characterization of 89 additional inter-protein pairs of the 214 total, a remarkable gain in informative protein-protein pairs.

### Crosslink Distances Distribution Reflects Native-Like Architecture

PhoX-mediated crosslinking imposes a characteristic spatial constraint, generally falling within a range of 7 to 25=Å (see Figure 3), owing to the lengths of lysine side chains (7=Å), backbone flexibility (6=Å), and the crosslinker molecule (∼7=Å). Accounting for structural flexibility and solvent accessibility, the experimentally verified upper limit is 33–35=Å (Bullock et al. 2018). Accordingly, we used this criterion to validate and map our crosslinking data onto available PDB structures for both plant species. All the distances measured in known complexes of spinach and *Arabidopsis* (8 and 5, respectively) are plotted in Figure 3A (Data S1). As an example of mapping, we show the So_P1 crosslinks mapped onto the PSII-LHCII supercomplex structure (PDB #3JCU, Figure 3B)

**Figure 3.**
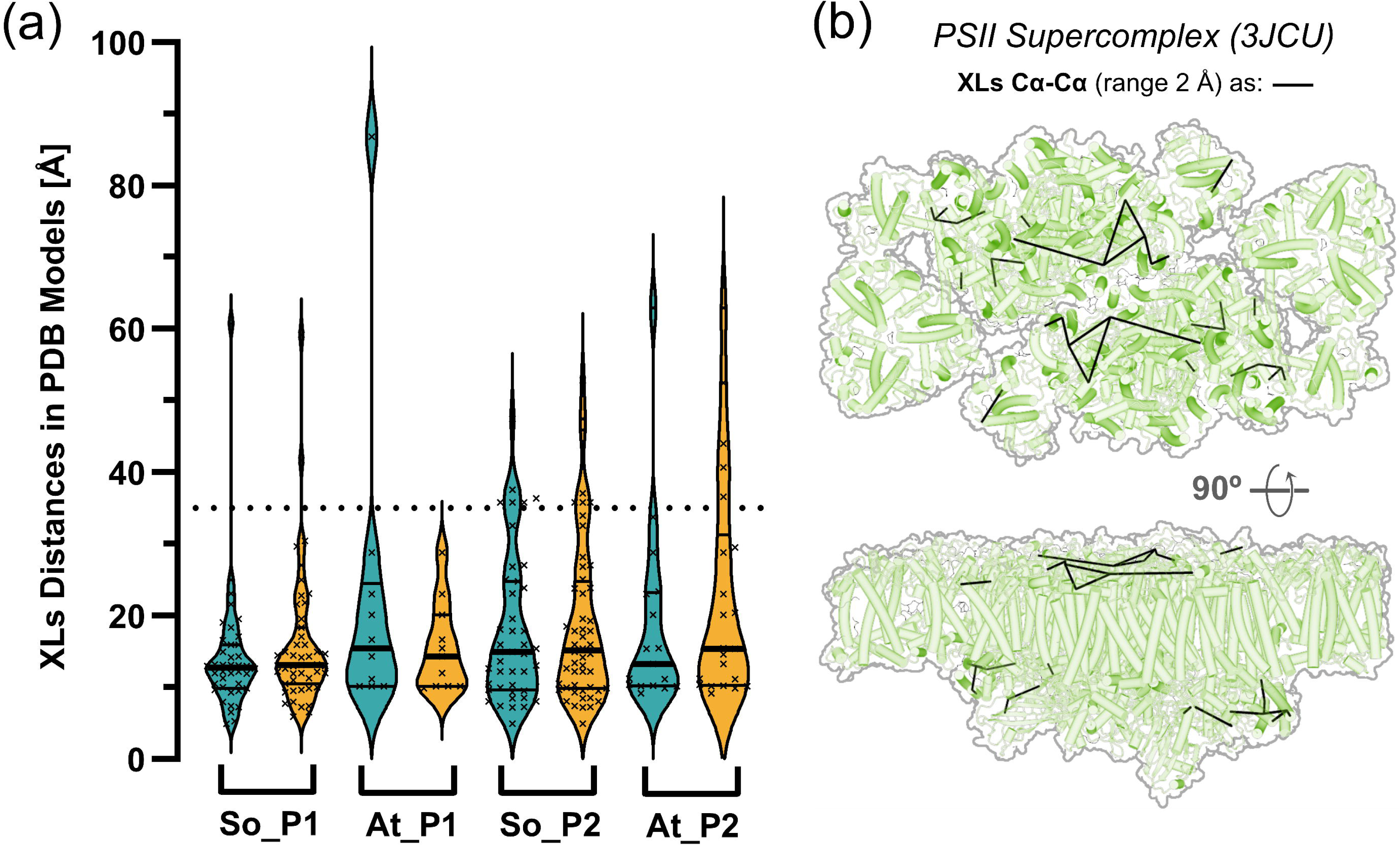
Structural validation of identified crosslinks by measuring distances in available PDB models of chloroplast protein complexes of spinach: 3JCU (PSII-LHCII), 6VON (Chloroplast ATP synthase), 7ZYV (Cytochrome B6f), 2WSC (PSI-LHCI), 7Q57 (GAPDH), 4V61 (Chloro-ribosome), 8RUC (RuBisCO), and 4RI2 (PsbS). In A. thaliana, five models were containing identified crosslinks: 7OUI (PSII-LHCII), 7WG5 (NDH-PSI-LHCI), 7UX9 (DDM1), 3RFY (Cyclophilin 38), and 5IU0 (RuBisCO). (a). Measured crosslink distances in PDB structures for A. thaliana thylakoids (At) or Spinach (So). Blue represents PhoX alone (1 mM and 2 mM as P1 and P2), and in yellow its combination with 100 mM TMPAC. When multiple copies of the same proteins where present in the PDB structure (i.e. for multimeric assemblies) only the shortest crosslink distance was considered. Each mapped crosslink distance is represented by an individual cross, and the horizontal line marks the maximum threshold distance of 35 Å. The central mark in the plot indicates the mean value along with its quartiles. (b) Example of PDB model of Photosystem II – LHCII supercomplex from spinach (3JCU) after fitting of all crosslinks (in black) found in So_P1 (2 out 3 replicates).

At 2 mM PhoX, longer-than-expected distances (>35 Å) were observed in *A. thaliana* and spinach (At_P2 and So_P2, Figure 3A). A smaller proportion of these outliers appeared in 1 mM PhoX-treated samples, particularly in spinach (So_P1), where most crosslinks remained within range. This may suggest that the treatment with 2 mM PhoX (P2) might not be the most appropriate, and 1 mM treatment (P1) may be preferable. Overall, over 97% of the detected crosslinks are within the expected distances for the 1 mM PhoX samples, and down to 85-90% for the 2 mM. This is consistent with the idea that higher crosslinker concentration may introduce longer or artefactual crosslinks (longer than the 35 Å cut-off, dashed line in Figure 3A) and that sample preservation can affect complex integrity. Importantly, the additional TMPAC treatment did not increase these average distances (Fig. 3), *i.e.,* it did not introduce artifacts. It is also worth noting that less than 10% of our data (84 out of 898 total peptide pairs for At and So) have structural matches to known experimental structures. This depends on both the relatively little number of known structures for the model organism *A. thaliana* compared to spinach, and on the higher yield of crosslinked sites on flexible and solvent-exposed domains (*e.g*. N-terminal loops of Light-harvesting complexes) as previously reported (Albanese, Tamara, Saracco, et al. 2020). This also underlines the complementarity between classical structural approaches and the use of XL-MS to uncover novel functional protein complexes *in situ*.

### A Scalable Framework for Mapping Functional Membrane Protein Networks Beyond Known Complexes

Once we validated our approach from a physiological and structural point of view, we focused on the interactome landscape, chasing novel protein pairs and unexplored interactions. Maps, for both species, were generated to retain crosslinks detected in at least two or three replicates. For completeness 1 and 2 mM datasets, with and without TMPAC, are visualized together (separate maps can be explored from Data S2). In spinach thylakoids (*Arabidopsis* is shown in Figure S4) the mapped protein network provides a complete overview of the reproducible protein-protein interactions identified in this study, along with curated protein names (see Table S3 for Uniprot and gene name mappings). The primary protein interaction networks involved PSII components, along with ATP synthase, Photosystem I (PSI), and, to a lesser extent, cytochrome b_6_f (Figure 4 and Figure S4). The identified crosslinked proteins could be categorized into three main groups: 1 - Well-characterized complexes of the photosynthetic electron transport chain (e.g., PSII, PSI and their LHC) with at least partial structural information available; 2 - Adjuvant or regulatory proteins, including LHC-like proteins (e.g., LIL, PsbS) and enzymes such as peptidyl-prolyl isomerases (PPIases); 3 - Uncharacterized or predicted proteins, such as plastid-lipid-associated proteins (PLAPs), for which little functional and structural information exists. This expanded network offers a more comprehensive view of chloroplast protein organization and provides new hypotheses for regulatory interactions within photosynthetic membranes. All maps are interactively accessible in the crosslinkviewer web application (https://crosslinkviewer.org, Data S2). A brief practical guide on how to interactively explore these protein-protein interaction networks is provided (Appendix 1).

**Figure 4.**
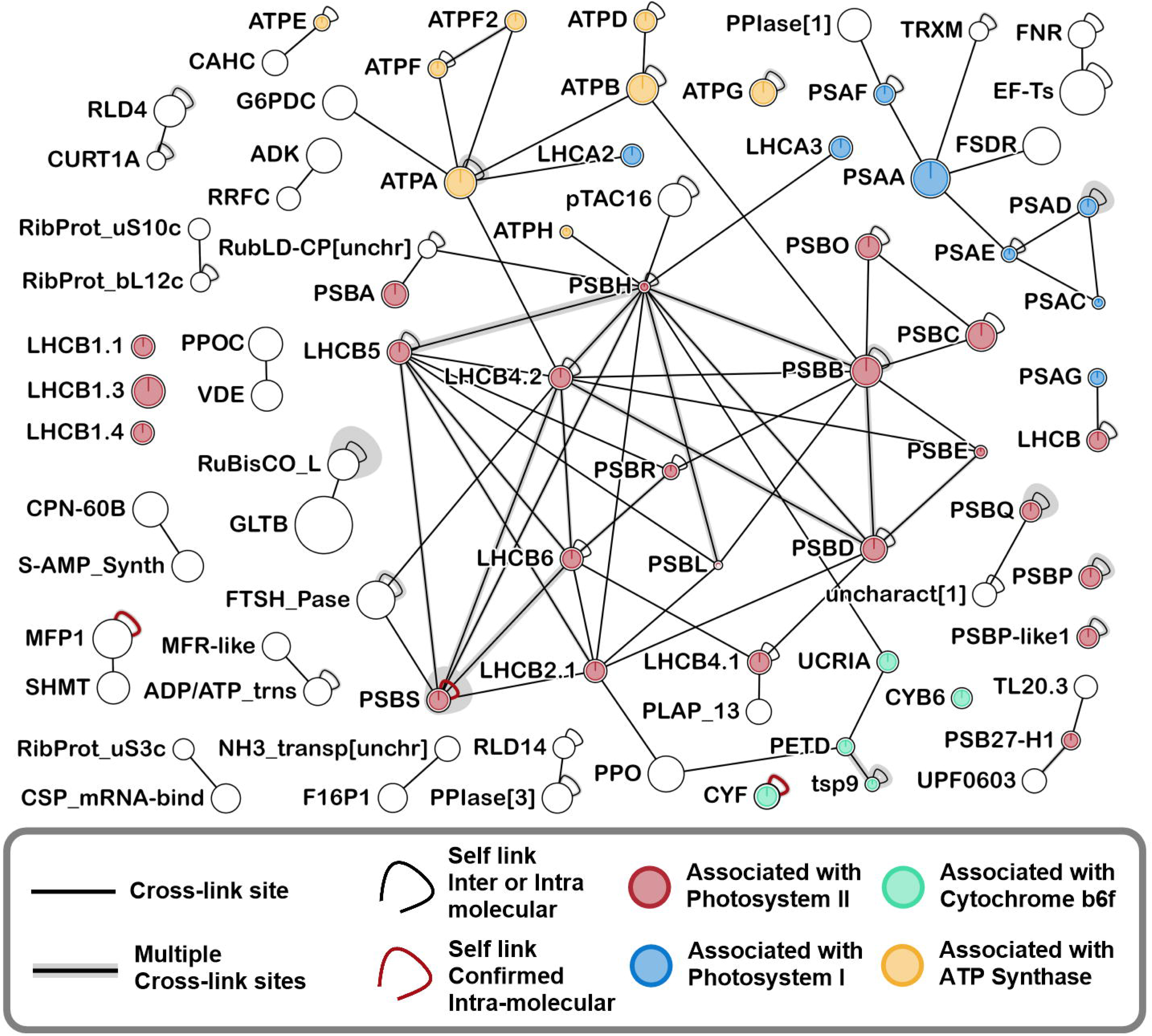
Interactome overview of thylakoids extracted from high-light Spinach acclimated leaves. (450 μmol photons m⁻² s⁻¹) crosslinked with 1 mM of PhoX crosslinker (with and without TMPAC). XLs present in at least 2 out of 3 replicas for each treatment are shown. Crosslink sites refers to dipeptides covalently bound; further details can be seen by opening the Supplementary Data in the web application https://crosslinkviewer.org. Color coding of the proteins are based on the information available about their known interaction with photosynthetic complexes, or during their assembly. Confirmed self-links indicate homomeric interactions and correspond to peptide pairs where identical residue positions are crosslinked or where peptides share overlapping sequences, configurations that are impossible within a single polypeptide chain (Lagerwaard et al. 2022). Due to the complexity in multi-interactions given by LHCB1 antenna proteins, these are not shown here (full view of the interactome in Data S1 and Appendix 1 for instructions).

Among the newly identified interactions, several point to previously unappreciated spatial associations within the thylakoid membrane. Some of the sub-networks may suggest the formation of structural-functional microdomains for regulating electron transport and PSII repair machinery (Lavergne and Joliot 1991; Kirchhoff et al. 2000). For instance, STR4 in Arabidopsis is also called TROL (Thylakoid Rhodanese-Like protein), and anchors ferredoxin–NADP⁺ reductase (FNR) to PSI, regulating the balance between linear and cyclic electron flow (Jurić et al. 2009). In contrast, CURT1A proteins are membrane curvature determinants enriched at grana margins and are essential for thylakoid architecture (Armbruster et al. 2013). The observed CURT1A–TROL association (Figure S4) therefore suggests a potential spatial coupling between membrane curvature domains and PSI-associated redox regulation. Additionally, the proximity between TLP18.3 (Sirpiö et al. 2007) and PSB27 (Chen et al. 2006), both implicated in PSII repair and assembly, is consistent with their participation in common PSII repair intermediates and supports a coordinated role in PSII maintenance in dark-adapted thylakoids.

### Integrative Structural Modeling Reveals Novel Functional Associations in Photosynthetic Complexes

Crosslinking MS provides distance constraints that are useful for predicting and validating structural models generated *in silico* (Graziadei and Rappsilber 2022; Burke et al. 2023). In this study we detected a relevant number of protein-protein interactions that belong to proteins with either a functional characterization but not a known structure, or neither of both (Figure 4 and Figure S4). Here, we highlight selected examples that address relevant questions in the field and open new avenues for elucidating the functions of previously uncharacterized proteins. Among several PSII interactors, we identified a previously described but functionally enigmatic protein named FIP (Figure 5c), whose knockdown mutant exhibits enhanced resistance to high light (Lopes et al. 2018). We observed FIP crosslinked to the PSII reaction center subunits PsbD and PsbE (Figure S4), an interaction not previously reported. Structural modeling with AlphaFold 3 (Abramson et al. 2024) supports this association, with all mapped distances falling within 35 Å. Given FIP’s proposed role in PSII turnover via association with FtsH proteases, our data suggest that FIP may act as a mediator of PSII disassembly or repair.

**Figure 5.**
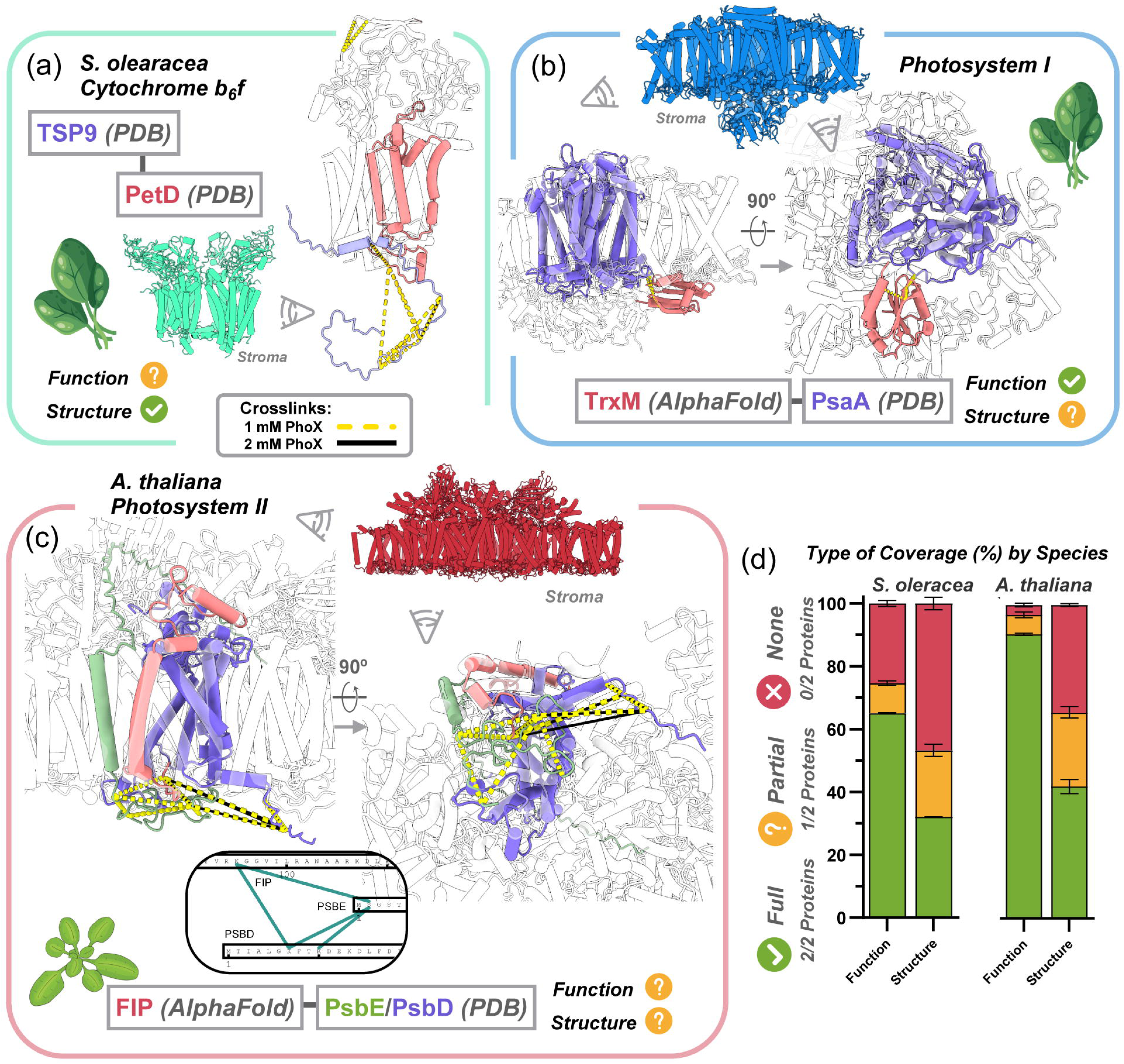
Structural and functional mapping of crosslinked protein interactions in *Spinacia oleracea* and *Arabidopsis thaliana.* Representative examples of crosslinks detected between photosynthetic complexes and associated proteins, shown in structural models of PSII, PSI, cytochrome b6f, and their interacting partners. Highlighted interactions include TSP9–PetD (a, So) and PsaA–TrxM (b, So), as well as FIP–PsbE/PsbD (c, At). Crosslinks are displayed as yellow dashed lines (1mM Phox) or black solid lines (2mM Phox). In panel c the network visualization of the inter-protein crosslinks is shown (see Data S1 and Appendix 1 for instructions). Bar plots (right) summarize the proportion of detected protein pairs with full (green), partial (orange), or no (red) annotation in terms of structural (PDB coverage) and functional (UniProt annotation) information across both species and PhoX concentrations (1 mM, 2 mM).

A common characteristic of uncovered interactors is the predominant targeting of intrinsically flexible or disordered regions. For instance, a thylakoid soluble phosphoprotein (TSP9) was reproducibly crosslinked to the PetD N-terminus of the cytochrome b_6_f, on the stromal side of the thylakoid membrane (Figure 5a). Previous work has proposed that TSP9 regulates light harvesting facilitating the dissociation of LHCII from PSII (Hansson et al. 2007; Fristedt et al. 2009). Here we show the reproducible detection of Tsp9 bound to PetD of the cytochrome b_6_f complex. This finding is in agreement with the presence of a rather small stretch of sequence of this subunit in a recent structure of cytochrome b_6_f (Sarewicz et al. 2023), and its possible absence in others due to a weaker interaction, which could be lost upon solubilization. Overlaying the predicted alphadfold2 structure (Jumper et al. 2021) of spinach TSP9 with the cryo-EM structure (PBD 7ZYV) of the cytochrome, we can show that 3 out of 4 inter-protein crosslinks with PetD are slightly longer than the threshold distance of 35 Å, namely between intrinsic disorder regions of TSP9 C-terminus. Our results confirm that TSP9 is part of the cytochrome b_6_f in dark adapted thylakoids.

Among the various modes of chloroplast electron transport, regulation often occurs at the level of Photosystem I (PSI), where pathways like cyclic electron flow (CEF) and the water–water cycle modulate energy production and redox balance. The latter contributes to ROS formation, as photoreduction at PSI generates superoxide, which is rapidly converted to H₂O₂ (Asada 1999). Detoxification relies on enzymes such as ascorbate peroxidases (APXs), catalases, and thioredoxins (TRXs) (Schürmann and Buchanan 2008; Courteille et al. 2013). Here, we observed the association of Thioredoxin M (TrxM) with the PSI core subunit PsaA (Figure 5b). The two interlinks found fit the *in silico* docked model. This proximity in dark-adapted thylakoids may suggest a localized redox relay, allowing TrxM to rapidly modulate PSI activity linked to ferredoxin (Dai et al. 2007).

Taken together, these findings illustrate how integrating crosslinking mass spectrometry with computational structural modeling and network analysis can uncover novel protein players within functional photosynthetic assemblies.

## Conclusions

This study establishes a robust and physiologically grounded approach to probe membrane protein interactions in functional thylakoid membranes using chemical crosslinking coupled to mass spectrometry. By demonstrating that photosynthetic activity is only partially affected during crosslinking, we show that structural proteomics can be applied directly to intact, light-responsive membranes — a critical advance for studying complex macromolecular assemblies *in situ*. We here introduced the use of TMPAC as an adjuvant molecule during the crosslinking reaction, initially motivated by physicochemical considerations rather than empirical optimization. As an amphiphilic cation, TMPAC likely reduces local charge repulsion and enhances crosslinker access to membrane-embedded or membrane-proximal proteins, without solubilizing the membrane. Importantly, physiological assays demonstrate preserved photosynthetic activity, and structural validation shows no increase in out-of-range crosslinks, indicating that improved yield does not come at the expense of structural integrity. Ultimately, we showed that TMPAC did not introduce a specific bias in the crosslinks identified, but rather diversifies them, thus representing a complementary approach that can increase by 20-40% the detected interactome landscape.

The resulting data reveal several reproducible interactions, and new contacts between core photosynthetic complexes, assembly and repair factors, and previously uncharacterized proteins. Structural validation across known 3D models confirms that most crosslinks fall within spatial constraints, and interactome maps illustrate dynamic connectivity across traditionally compartmentalized thylakoid domains.

Overall, we propose XL-MS as a powerful tool to bridge functional and structural biology in plants, enabling the systematic exploration of protein networks under near-physiological conditions. Our workflow is rapid, reproducible, and adaptable, offering a practical framework for investigating plastid biogenesis, energy metabolism, and stress responses in photosynthetic systems. Looking ahead, the integration of this approach with quantitative crosslinking, cryo-EM, and machine-learning-based structural modeling will provide unprecedented insight into the organization and regulation of energy-converting organelles.

## Materials and Methods

### Thylakoid Membrane Extraction

Wild type plants of *Arabidopsis thaliana* Col-0 (At) and *Spinacia oleracea* (So) cultivar ‘Paris HF1’ (Vilmorin), were grown at 22°C with a 12/12-hour light/dark cycle and a light intensity of 100 μmol m⁻² s⁻¹ for about six weeks. To perform “High-Light” treatment, plants were transferred to a growing chamber with an illumination of 450 μmol photons m⁻² s⁻¹ for 10 days. Leaves were harvested in a cold chamber at 4 °C under dim green light at the end of the night cycle. The leaf tissues were ground in a blender using a base buffer containing 20 mM HEPES pH 8, 5 mM MgCl2, 22 mM KCl, and 10 mM NaCl, with an additional 0.3 M of sucrose. For every 50 g of *A. thaliana* leaves (At), 900 mL of buffer was used, and 100 g for spinach leaves (So) for the same buffer volume. The plant material was blended in 15 separated pulses of about 1 second each. The resulting paste was filtered through a quadruple-layered cheesecloth and centrifuged at 3000g for 5 min. The pellet was resuspended in the same base buffer supplemented with 0.1 M sorbitol. The chlorophyll content was measured by diluting thylakoids in 80% acetone, as described by Porra et al. (2018). The samples were stored adding 1 M betaine (CAS:107-43-7) in the same resuspension buffer (in the dark), then quickly frozen in liquid nitrogen and eventually stored at −70 °C. For samples At_P1 and So_P1 and samples prepared for confocal imaging, thylakoids were used directly for the experiments, without freezing.

### Confocal Microscopy Imaging

Imaging was performed using a Zeiss LSM 900 confocal microscope equipped with ZEN software (version 3.0, Blue edition) with minor modification from previous set-up (Giustini et al. 2025). A 63x/1.4 M27 objective lens was used in confocal mode with an Airyscan 2 detector to ensure fast acquisition without compromising sensitivity, aided by a collimation optic. Excitation for chlorophyll fluorescence was provided by a 405 nm blue laser, and emission was collected between 650 and 700 nm. Nile Red visualization involved 488 nm excitation with fluorescence collected between 557 and 592 nm.

### Determination of the Maximum Quantum yield of PSII and Thylakoid Photosynthetic Activity

To evaluate the effects of TMPAC on thylakoid membranes during photochemistry, we measured the maximum quantum yield of PSII (Fv/Fm) and the photosynthetic electron transport rate to assess photosynthetic efficiency. Photosynthetic electron flow was examined under several conditions: only thylakoids (control), 100 mM TMPAC, 1 mM PhoX, 2 mM PhoX, and combinations of PhoX with TMPAC.

Chlorophyll fluorescence was quantified using a Speedzen III fluorescence imaging setup (JBeam Bio, France). Maximum fluorescence (Fm) was determined by saturating red pulses (520 nm, 250 ms, 3000 µmol photons m^-2^ s^-1^) and 10 µs detection pulses of blue light (470 nm). Minimum fluorescence in the dark was determined by using the same detection pulses in the absence of saturating pulses (F0) The maximum quantum yield of PSII was calculated as Fv/Fm=(Fm-F0)/Fm (Maxwell and Johnson 2000). The electron transport rate (ETR) was measured using the same setup and increasing the intensity of a continuous light at different light intensities after every 5 minutes of illumination for a total measurement time of 40 minutes (including a 5-minute dark period at the end of the light phase to evaluate sample recovery).

### Experimental Setup for Protein Crosslinking

Thylakoid solutions were treated with addition of 1 or 2mM PhoX (100 mM stock solution in anhydrous DMSO) or in combination with 100 mM TMPAC (1M stock solution in water). Detailed conditions are in Table S1. Four experimental conditions were set, the combination of At and So with PhoX 1 mM (P1) and 2 mM (P2), denominated as At_P1, At_P2, So_P1, So_P2. The reaction mix was kept in the dark at room temperature, and it was stopped after 30 minutes with 50 mM NH_3_. An extra step of thylakoid washing was performed in At_P1 and So_P1 to remove soluble proteins that were not covalently crosslinked (*i.e*. unspecifically interacting). Crosslinked thylakoids were washed twice by centrifuging at 3000×g for 5 min and resolubilized in 1 mL of base buffer before final resuspension at the final concentration of 2mg/mL of Total Chlorophyll. Another crucial aspect is that P1 treatment of crosslinking was performed the same day of extraction before betaine snap freezing.

Total protein was precipitated by adding ice-cold acetone in steps of 50%, 50%, 100%, 100%, and 200% of the initial solution volume, with shaking at 1000 rpm for 5 minutes between additions. Samples were stored overnight at −20°C and centrifuged at 22,000×g for 30 minutes at 4°C. The dry pellet was then resuspended in 300 µL of 1% sodium deoxycholate (SDC) and 100 mM tris pH 8 using mechanical assistance (2 glass beads of 4 mm in a 2 mL tube at 1000 rpm for 2 hours).

### Crosslinking Mass Spectrometry Sample Preparation

Digestion was carried out overnight at 37°C using trypsin from bovine pancreas (Merck) at a 1:20 enzyme/protein ratio. Trypsin activity was stopped by heating at 90°C for 10 minutes. To dephosphorylate endogenous peptides, the digested solution was supplemented with 25 mM MgCl_2_, 50 mM KCl, and centrifuged at maximum speed for 5 minutes. The supernatant was separated, and the pellet was resuspended with 100 µL of 100 mM tris (pH 8). This solution underwent an additional centrifugation step to extract any remaining peptides, which were combined with the previous solution. Calf intestinal phosphatase (Quick CIP, NEB) at a concentration of 10 units/mL was added to the recovered supernatant and incubated at 37°C for 8-10 hours.

Precipitation of residual SDC was induced by adjusting pH to 3 with formic acid. The solutions were then centrifuged to remove debris. Peptides were desalted using an Oasis HLB 96-well plate according to manufacturer’s protocol (Waters Corporation). Cross-linked peptides were enriched using the MagReSyn® Zr-IMAC protocol (Resyn Biosciences) with minor adjustments compared to the manufacturer protocol, namely we used a beads:peptide ratio of 1:4 we increased the glycolic acid concentration to 1M in the binding buffer, and NH_3_ to 5% in the elution buffer. Beads were recycled by repeating the entire protocol with the same nanoparticles for each sample and then joining the final eluates, as it was shown to increase the recovery (Bortel et al. 2024).

### Mass Spectrometry Analysis

Phospho-enriched peptides were processed in two different NanoLC-MS/MS configurations: a Dionex UltiMate 3000 RSLCnano coupled to an Orbitrap Exploris 480 through an FAIMS Pro™ interface (CV 55-60-75) (Thermo Fisher Scientific) and a UHPLC Vanquish Neo coupled to an Orbitrap Ascend Tribid mass spectrometer (Thermo Fisher Scientific). Peptides were separated on an Aurora Ultimate C18 column (IonOpticks) using a 120-minute gradient from 2.5% to 41% acetonitrile in 0.1% formic acid at 300 nL/min. Full scan MS spectra covered 400 to 1600 m/z. MS2 ions were fragmented with normalized collision energy of 21-28-36 for the Orbitrap Exploris and 21-29-37 for the Ascend. Ions with less than 2+ charge were excluded.

### Evaluation of Monolink Abundance

Raw files were analyzed with FragPipe 22.0 (Kong et al. 2017) for modifications on lysine residues from PhoX. Isotopic masses considered were those of PhoX reacting with water (227.983 Da), ammonia (226.998 Da), and side derivatives (PhoX anhydride: 209.972 Da and PhoX-acetamide: 338.067 Da). The search was repeated with the identified proteins, this time seeking to identify semi-tryptic cleavage sites, which enabled the detection of amino terminal residues resulting from post-translational cleavages and mature proteins after cleavage of the transit peptide.

### Analysis of Crosslinked Peptides

For Crosslinks search data analysis was conducted using XlinkX Node in Proteome Discoverer 3.0 (Thermo Fisher Scientific) to identify cross-linked (XL) peptides with a minimum score cut-off of 40 (Klykov et al. 2018). The 20% most abundant identified proteins in the mono-link analysis (FragPipe output) were implemented as a database along with the inclusion of proteoforms with non-tryptic N-terminal cleavages. The proteoforms were obtained after including non-tryptic cleavage in FragPipe settings.

In order to conduct the study, only those cross-links that appeared in at least two out of three replicates (2 out 3) or all three replicates (3 out 3) for each condition were considered. Data filtering and formatting were conducted using an R script, which was subsequently employed to reshape the data into the input for xiNET, an online cross-link viewer developed by the Rappsilber Laboratory (Combe et al. 2015). Crosslinked were mapped onto PDB structures using the ChimeraX plugin XMAS (Lagerwaard et al. 2022)

### Custom R-Based Pipeline for Re-Annotation and Visualization of Cross-Linked Peptides in XiNET

To analyze cross-linked peptide data generated by Proteome Discoverer (PD), we developed a custom R-based pipeline for re-annotation and visualization of XL-peptides in xiNET. The analysis was performed using R version 4.2.1, incorporating packages such as tidyverse, dplyr, Biostrings, data.table, stringr, and seqinr. Input data consisted of crosslink spectrum matches (CSM) exported from PD and a reference protein sequence database.

The pipeline organized data according to MS run identifiers and classified them by experimental condition. Data pre-processing included filtering out cross-links that did not meet a user-defined minimum XlinkX score threshold (for this analysis 40) and grouping them based on a minimum number of repeated cross-link hits across replicates (“2 out of 3” and “3 out of 3” replicates). Remaining data were consolidated into unique cross-links, defined by sequence and site of crosslinking, retaining the highest score, the number of spectral matches, and the cumulative precursor abundances from aggregated CSMs. Cross-links were annotated with peptide sequences, linking positions, and associated scores, and formatted for visualization in xiNET.

Protein sequence re-annotation was performed by mapping peptides to a FASTA database, refining annotations originally assigned by PD This step aims to eliminate redundant proteoforms, essential for finding XLs with N-terminal residues after transient peptide cleavage. Furthermore, the pipeline generated all possible protein-pair combinations by aligning peptides to the updated database. This step is applied to distinguish between interactions that might occur between multiple proteins sharing the same peptide sequences due to high homology (e.g. light harvesting complex proteins), from protein-specific interactions. The complete R scripts used to handle the data are available at the GitHub repository https://github.com/LPM-team1/N-TRIMS

## Author Contribution

Conceptualization – NF, MF, GF, PA

Data Acquisition – NF, CG, PA

Data Analysis – NF, PA

Supervision – MF, GF, PA

Manuscript Writing - NF, PA

Editing - NF, MF, GF, PA

## Acknowledgements

We would like to thank all the members of the LPM and EDyP laboratories for their support and assistance, especially A.M. Hesse, L. Belmudes, J. Novion-Ducassou, S. Kieffer-Jaquinod, S. Miesch-Frémyand and M. Menneteau for their invaluable technical support. This research project was funded by the 2022 ANR PRC program (PHOTO_DYN - 22-CE20-0015-01). P.A. acknowledges funding from the European Union’s Horizon 2020 research and innovation program under the Marie Skłodowska-Curie grant agreement No 101066400 - PhotoLINK. GF acknowledged funding from the European Research Council ERC (Chloro-Mito; grant no. 833184) and the EIC PlanktON project (grant no. 101099192). Proteomic experiments were partially supported by Agence Nationale de la Recherche under projects ProFI (Proteomics French Infrastructure, ANR-10-INBS-08; ANR-24-INBS-0015). This work was also supported by ANR-17-EURE-0003 and GRAL and/or ANR-10-LABX-49-01

## Data availability

All the raw data files are provided as Source Data to the corresponding figures. Code and scripts used to process and analyze fasta files and XL-MS data are accessible at the LPM GitHub repository https://github.com/LPM-team1 (N-TRIMS and xiNET-Mapping-Algorithm). All MS raw data files, including the search results, are deposited to the ProteomeXchange Consortium via the PRIDE partner repository (dataset identifier PXD068425). Source Data for Figures 1,3,4 and 5 are deposited and accessible via Zenodo (https://zenodo.org/records/17308311)

*For reviewers, MS data can be accessed at the following address:* https://www.ebi.ac.uk/pride/login.

*Username: Password: JUWG6a8fc3Pm*

---

**Figure S1.**
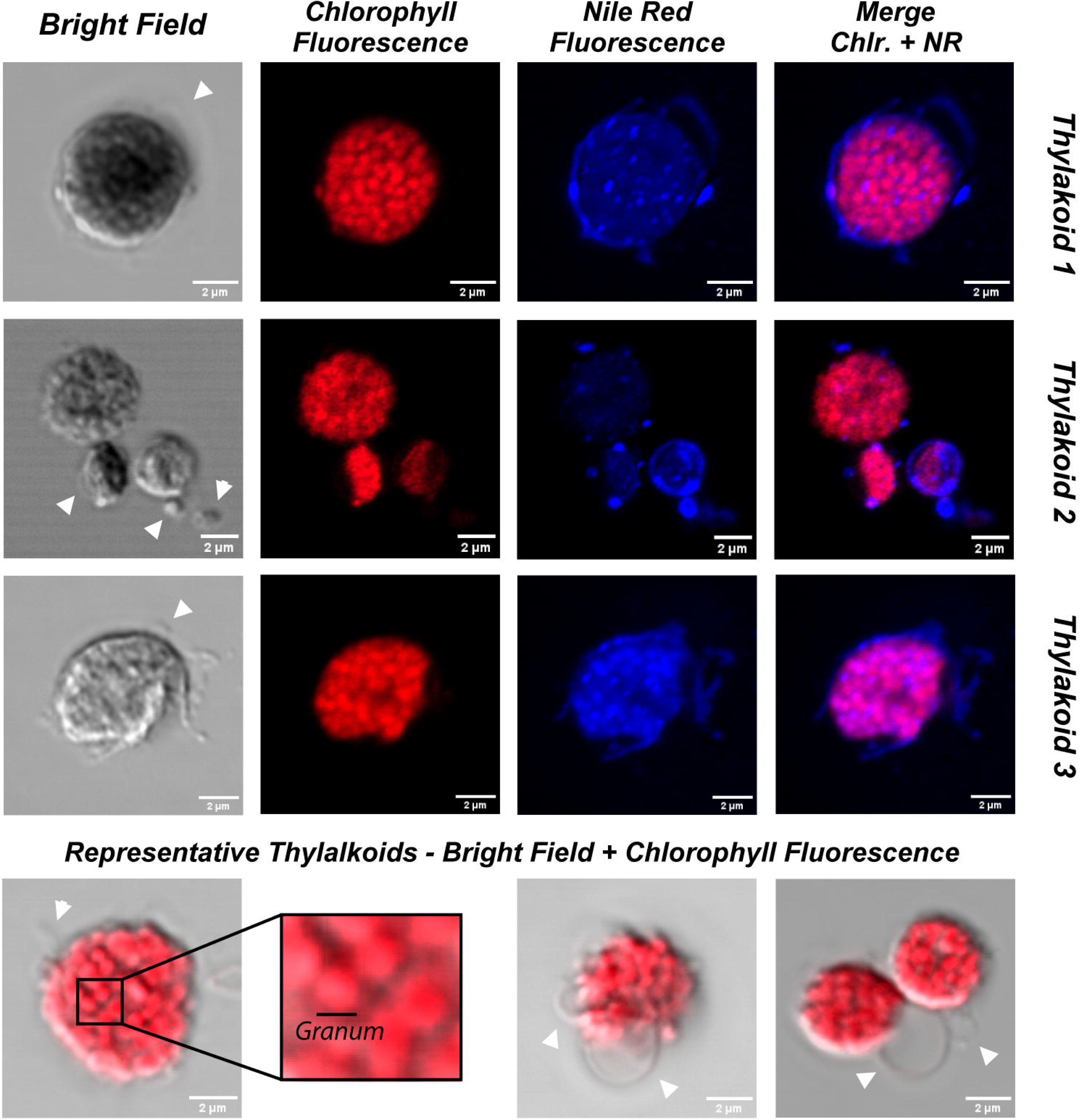
Confocal imaging of isolated thylakoid preparations. Representative bright-field and confocal fluorescence images showing chlorophyll autofluorescence (red) and Nile Red staining of membrane lipids (blue) in purified thylakoid membranes. Merged images highlight the overlap of chlorophyll and lipid fluorescence. Arrowheads indicate smaller membrane fragments visible in some preparations. Scale bars = 2 µm.

**Figure S2.**
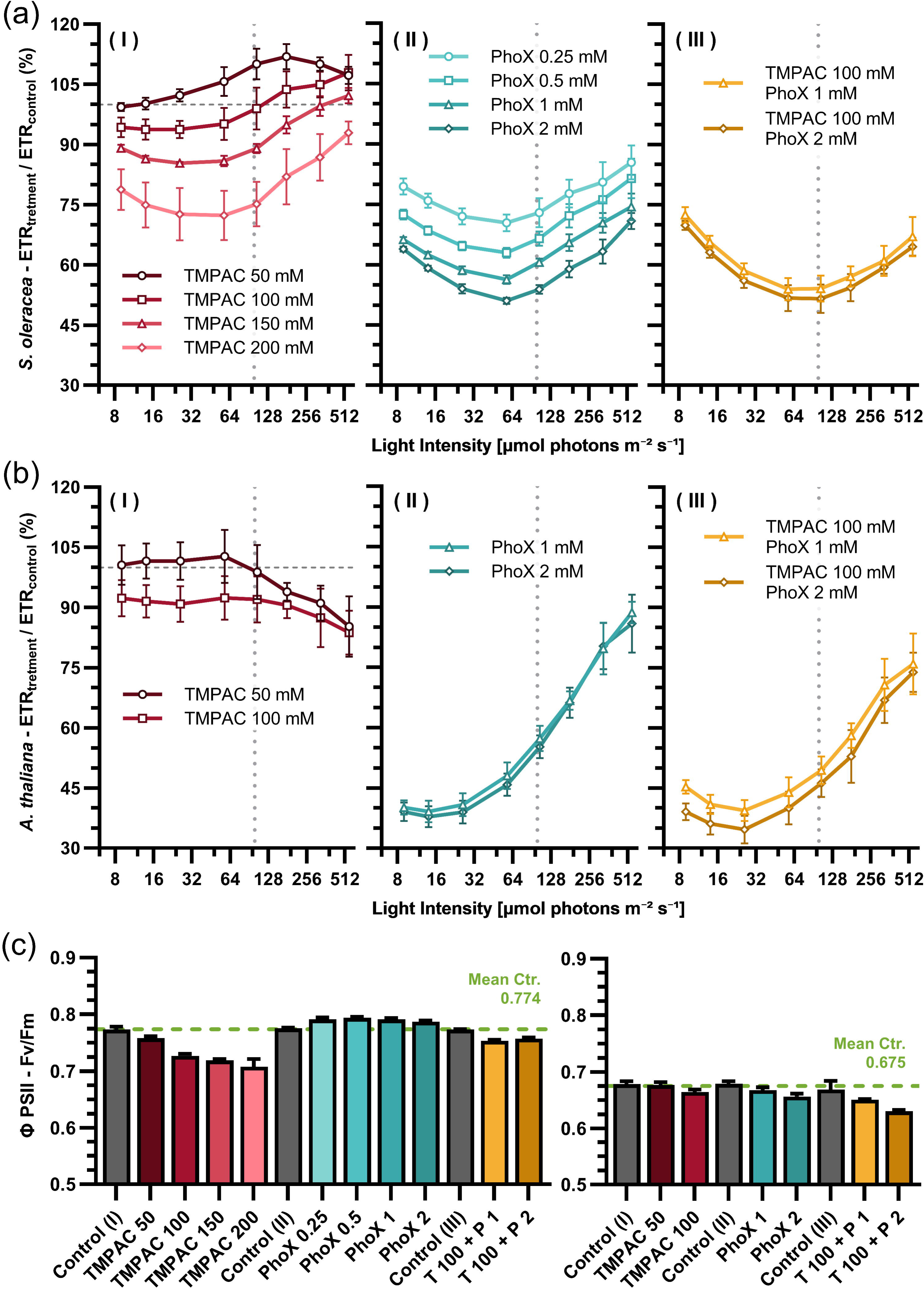
Photosynthetic performance in response to increasing TMPAC and Phox crosslinker concentrations. (a) Electron transport rate (ETR), expressed as percentage of the respective control, measured in spinach thylakoids across light intensities from 8 to 512 µmol photons m⁻² s⁻¹. (I) TMPAC (50–200 mM); (II) Phox (0.25–2 mM); (III) TMPAC 100 mM combined with Phox (1 or 2 mM). Roman numerals (I–III) indicate measurements obtained from the same thylakoid batch and recorded simultaneously. The vertical dashed line marks 100 µmol photons m⁻² s⁻¹ (plant grow condition) and horizontal dashed lines marks the reference control value (100%). Error bars represent SD (n > 3). (b) ETR measured in A. thaliana thylakoids under selected TMPAC (50, 100 mM), Phox (1, 2 mM), and combined treatments, across the same light range. Labels are the same as in panel (a). (c) Maximum PSII quantum yield (Fv/Fm, dark-adapted) determined for the samples shown in panel (a) and (b), spinach and A. thaliana, respectively. The combined treatment of TMPAC 100 mM with PhoX is indicated as T100 + P1 or T100 + P2, corresponding to PhoX at 1 mM or 2 mM, respectively. Green dashed lines indicate the mean control value for each species. Error bars represent SD (n > 3).

**Figure S3.**
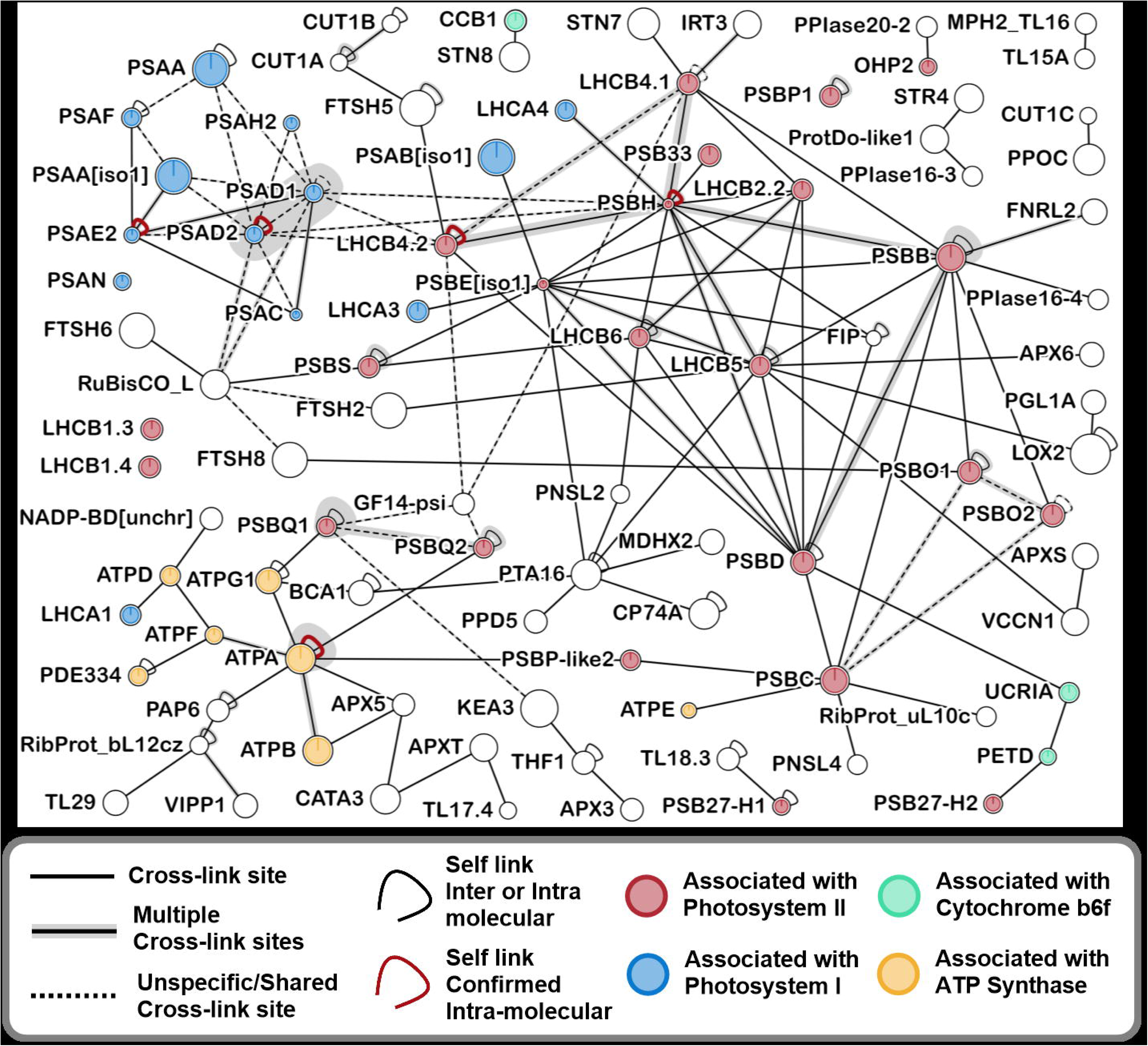
Amino acid class distribution in the ±3 residue window surrounding PhoX crosslinking sites under different experimental conditions. Bar plots show the relative frequency (%) of amino acid classes neighboring the crosslink position for peptides common to both conditions (left column), peptides unique to PhoX-only experiments (middle column), and peptides unique to TMPAC+PhoX experiments (right column). Data are shown for *Spinacia oleracea* (So) and *Arabidopsis thaliana* (At) samples analyzed at two PhoX concentrations: 1 mM (P1) and 2 mM (P2). The total number of crosslinked peptides (XL pept) and corresponding single peptides contributing to each analysis are indicated within each panel. Amino acid classes are defined as hydrophobic (Hf), polar (Po), special cases (Sc; G, P, C), positively charged (C+; K, R, H), negatively charged (A−; D, E), aromatic (Ar; F, Y, W), N-terminal (Nt), and C-terminal (Ct). The dashed boxes report the ratio between positively and negatively charged residues (Cation/Anion) for each condition

**Figure S4.**
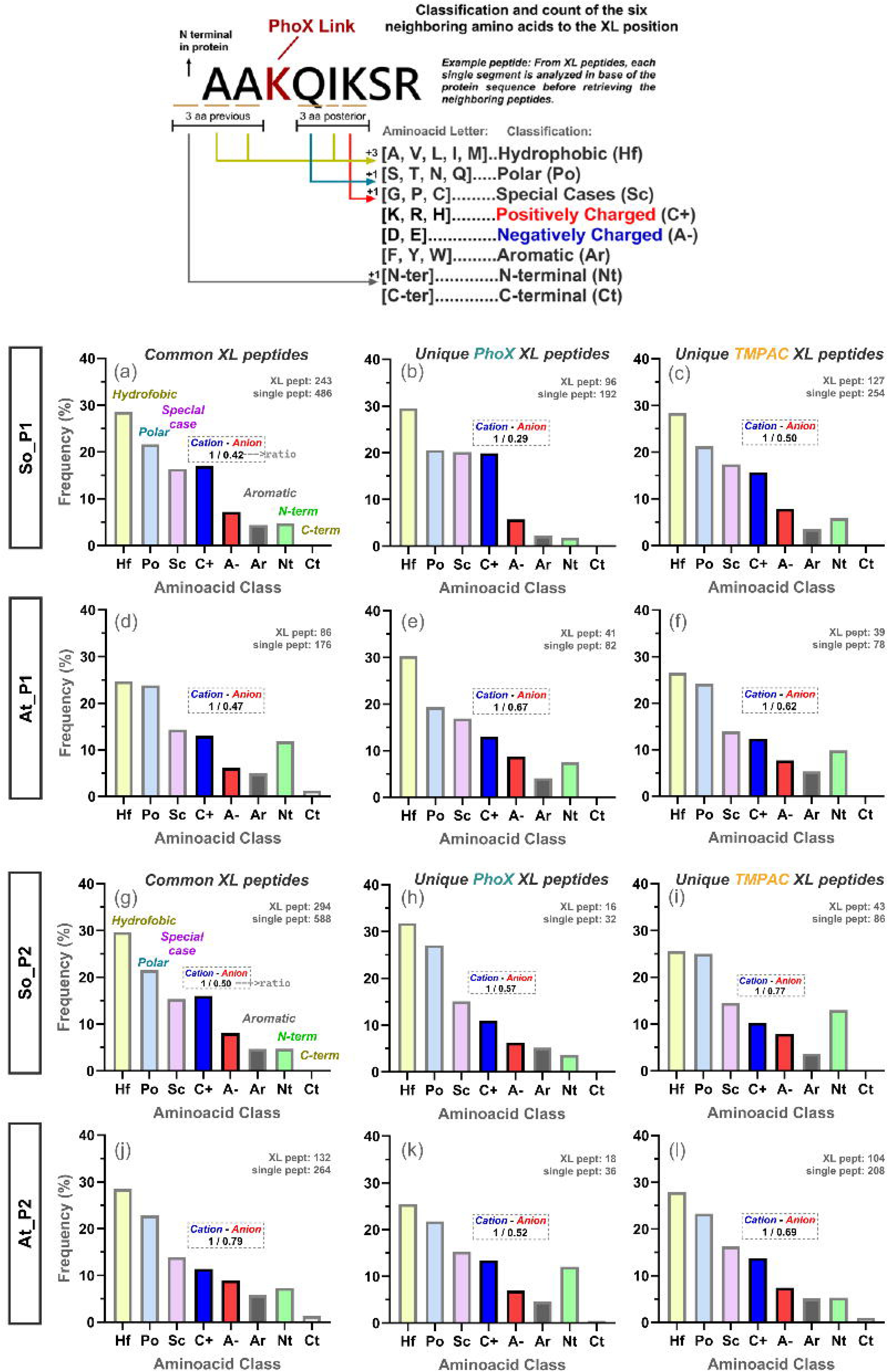
Interactome overview of thylakoids extracted from low-light acclimated *Arabidopsis thaliana* leaves. (100 μmol photons m⁻² s⁻¹) crosslinked with 1 mM of PhoX (with and without TMPAC). XLs present in at least 2 out of 3 replicas for each treatment are shown. Crosslink sites refers to dipeptides covalently bound; further details can be seen by opening the supplementary data S2 in the web application https://crosslinkviewer.org. Color coding of the proteins is based on the information available about their known interaction with photosynthetic complexes, or during their assembling. Confirmed self-links indicate homomeric interactions and correspond to peptide pairs where identical residue positions are crosslinked or where peptides share overlapping sequences, configurations that are impossible within a single polypeptide chain (Lagerwaard et al. 2022). Due to the complexity of multi-interactions in LHCB antenna proteins, we highlighted them in grey to allow for a tidier visualization (full view of the interactome in Data S1)

**Table S1.** Experimental conditions for crosslinking of thylakoid membranes.

Growth light acclimation, crosslinker concentrations, TMPAC treatments, and sample-specific parameters for *Arabidopsis thaliana* (At) and *Spinacia oleracea* (So) thylakoid preparations. Values include maximum quantum yield of PSII (Fv/Fm, error <0.8%), total chlorophyll concentration, reaction volume, washing steps, protein-to-chlorophyll ratios, and total digested protein amounts.

**Table S2.** Annotated XL-MS data output across experimental conditions.

Cross-linked peptide identifications from thylakoid samples of *Spinacia oleracea* (So) and *Arabidopsis thaliana* (At) under control (PhoX only) or TMPAC-treated conditions. Each sheet corresponds to one experimental setup: So_1mM_ctrl, So_1mM_TMPAC, So_2mM_ctrl, So_2mM_TMPAC, At_1mM_ctrl, At_1mM_TMPAC, At_2mM_ctrl, and At_2mM_TMPAC. Listing peptide sequences, crosslinking residues, protein assignments, replicate occurrence, and quantitative parameters (scores, spectral counts, precursor abundances). Additional annotation columns were appended to the original Proteome Discoverer software output table to provide functional information retrieved automatically from UniProt, including gene names, ORF identifiers, protein descriptions, annotated functions, GO/localization terms, and PDB codes when available.

**Table S3.** Annotated protein names and sequences.

Annotated custom protein names, gene names when available, Uniprot IDs and full-length amino acid sequences of all proteins included in the analyses presented in this study.

**Table S4.** Structural mapping of identified crosslinks.

Distance measurements for mapped crosslinks in available PDB structures of chloroplast complexes. Distances are reported as shortest Cα–Cα measurements, categorized per PDB entry and experimental condition, with indication of in-range (<35 Å) and out-of-range (>35 Å) values.

**Appendix 1.** A brief practical guide on how to interactively explore these protein-protein interaction networks is provided

**Data S1.** Comprehensive compilation of structural complexes, crosslink assignments, and calculated crosslink distances (0 Å and 1 Å margins) across samples and treatments, including filtered crosslinks (FXL), model annotations, and summary tables used for violin plot analyses.

**Data S2.** Complete dataset, including the full interaction table and corresponding FASTA sequences, formatted for upload to crosslinkviewer.com

