## Appendix 1 for "Deciphering Photosynthetic Protein Networks: A Crosslinking-MS Strategy for Studying Functional Thylakoid Membranes"

#### Overview of Raw Data and Visualization Resources

This document serves as a guide to the structure, content, and usage of the datasets and visualization tools associated with the manuscript on crosslink-based proteomic analysis. It explains where to find the raw data files used in the study and how to visualize interactomes using online tools.

#### Raw Data Documents

Within the folder **Crosslink_Peptides** (*Extended_Data_1\...\Crosslink_Peptides*), you will find the complete list of crosslinks accounted for in the thesis analyses. These tables (*Frequent_Crosslinks_(In at least 2 rep)...*) include only crosslinks that have passed the following minimum identification parameters:

- False Discovery Rate (FDR) threshold of 1%
- Proteome Discoverer score > 40
- Presence in at least 2 MS replicates out of the 3 datasets.

Key Columns within Each File

| Column | Description |
| --- | --- |
| **RawFile_Name** | Name of the MS run |
| **Experimental group** | Experimental condition (e.g., PhoX/TMPAC treatment or NPQ crosslinking time window) |
| **Sequence_A / Sequence_B** | Peptide sequences of crosslinked partners |
| **Position_A / Position_B** | Positions of the crosslinker attachment sites |
| **Protein_A / Protein_B** | Proteins corresponding to each peptide |
| **ID.Uniprot_Prot_A / ID.Uniprot_Prot_B** | UniProt identifiers of each protein |

### 🧬 Guide for Visualization of Interactomes

#### 1. Accessing the Viewer

To visualize interactomes, open the following link in any modern web browser:
🔗 https://crosslinkviewer.org/upload.php

You will see the upload window with several input fields for your files.

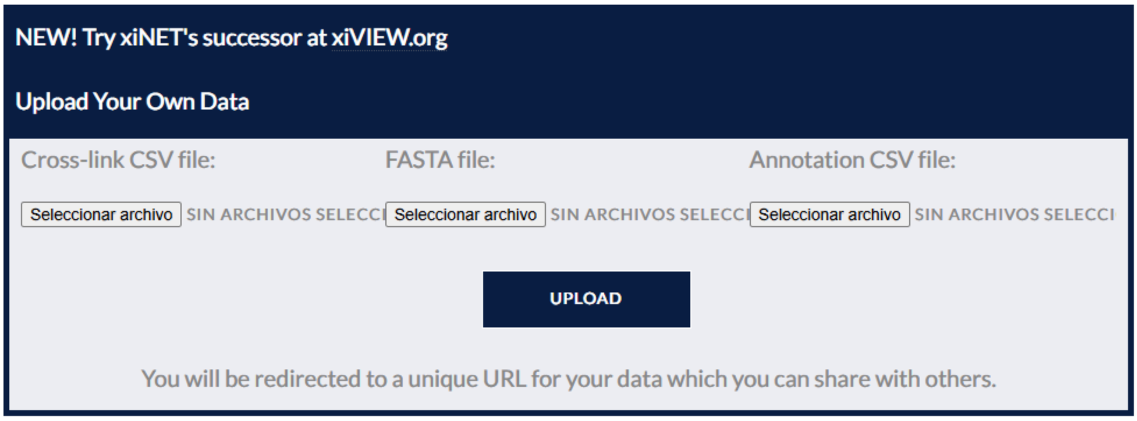


#### 2. Selecting the Files to Upload

You will find two main folders containing the interactome datasets:

- **At_P1 / At_P1_TMPAC / At_P2 / At_P2_TMPAC** → *Arabidopsis* (treated with 1 or 2 mM PhoX, and with 100 mM TMPAC respectively)
- **So_P1 / So_P1_TMPAC / So_P2 / So_P2_TMPAC** → *Spinach* (treated with 1 or 2 mM PhoX, and with 100 mM TMPAC respectively)

All conditions are filtered <40 in Proteome Discoverer Score and are found at least in 2 out of 3 replicates per treatment.

#### 3. Uploading the Files

Once you have selected the condition and the corresponding .xinet or .csv file, follow these steps:

1. **In the CrosslinkViewer page**, locate the field labeled:
   ➤ *"Cross-link CSV file"*
   Upload your .csv file containing the crosslink data here.

   *Example:* ***So_P1_TMPAC.CSV***

2. Depending on whether you are working with **Arabidopsis** or **Spinach**, upload the additional required files found at the **beginning of the folder**:

   - **FASTA file** → contains the protein sequences.

     *Example:* ***Fig4_So.FASTA***

- **Annotation CSV file** → contains the protein identifiers and descriptions.

  *Example:* ***Fig4_So_Annotations.CSV***

✅ *Make sure the correct FASTA and annotation files match the organism of your dataset.*

##### 4. Example Datasets

Here are example links showing preloaded interactomes for each organism:

- **Spinach (So_P1_TMPAC):** https://crosslinkviewer.org/uploaded.php?uid=8b09038e16505417ce1f3e8b8d131844042c85c6
- **Arabidopsis (At_P1_TMPAC):** https://crosslinkviewer.org/uploaded.php?uid=f9b5899923d8ad70342c0c6acdd3a986fee61d72

##### 5. Navigating the Interactome

Once your interactome is loaded, you can explore it using the following controls:

🖱️ **Right-click** → Expand the sequence of a protein.
🖱️ **Shift + Right-click** → Further expand the protein sequence (multiple times).
🖱️ **Right-click + Drag** → Move (reposition) protein globes within the visualization space.
🖱️ **Left-click** → Hides crosslinks from a protein globe.
