## Supplementary material for "Deciphering Photosynthetic Protein Networks: A Crosslinking-MS Strategy for Studying Functional Thylakoid Membranes": Source Data S1: README.txt

In this folder you will find: "Distances in Complexes from Samples"
1st Foglio Excel: List of complexes that contains the protein comprised in each peptide from a cross-link.

2nd Foglio Excel: List of complexes as a resume of 1st Foglio Excel. Entry complex tag, variants of the model by entry tags. Times that the complex was detected with an inter or intra XL. How are named the ChimeraX (XMAS) file with all posible distances fitted into the model, and the name of the subset file by 1 Å margin. Content inside the PDB model, and notes related whether is already encomprised in another bigger model, or it is not interesting (Keratine).

3rd Foglio Excel and so on: The distances obtained by PDB model with a lenght restriction of 0 Å margin. As a rule, one distance per crosslink in each model, avoiding over-representation in the violin plots. All XLs have the same weight, idependintly if the proteins where they are found are multimeric or XLs have multiples matches.

Last Foglio Excel: The table with all XL distances (0 Å margin) per sample and treatment. # Violin plots comes from here! 

- Then: Folder with the different samples names.
- Inside: 
- FXL + Experiment (.xlsx) (FXL - Filtered Cross-links) Either all XLs, XLs in 0 mM TMPAC, or XLs in 100 mM TMPAC.
- ShortSubset + Entry PDB + Condition (.xlsx) XL distances with 0 Å margin.

-- Then: Files folder
-- Inside: 
-- Entry PDB + Condition (.xlsx) Either all posible distances from all XLs.
-- Entry PDB + Condition + 1A_Short (.xlsx) XL distances with 1 Å margin. Do I use it for something? No. :D

#---------------------------------
#  R code to get XLs distances
#---------------------------------

library(tidyverse)
library(dplyr)
library(writexl)

# Set working directory and define folder paths
setwd('X:/NICOLAS/Documents/Trabajo/Doctorado-CEA/Bureau_De_Trabajo/R_Coding/R_Analisys/Distance_Clean')
Folder1 <- 'Paper_Dec24/So_P2_2o3'
Folder2 <- '/Files'
File_comp <- paste0(Folder1, Folder2)

# Define variables
#PDB_Model <- c('7OUI', '7WG5')
PDB_Model <- c('2WSC','3JCU','4V61','6VON','7ZYV','8RUC','7Q57','2PUK') #Spinach 2P // PDB_Model <- c('2WSC','3JCU','4RI2','4V61','6VON','7ZYV','8RUC') Spinach 1P // PDB_Model <- c('7OUI','5IU0','7UX9','7WG5','3RFY') Arabidopsis
Treat_Name <- c('_(0mM)', '_(100mM)')
Subset_Name <- '_1A_Short'
Do_Subset <- 'No'
Min_Distance <- 0


my_comp <- data.frame()

# Only process files if Do_Subset is 'No'
if (Do_Subset == 'No') {
  for (Z in 1:length(PDB_Model)) {  
    for (X in 1:length(Treat_Name)) { 
      # Construct file name and path
      Name_comp <- paste0(PDB_Model[Z], Treat_Name[X], '.tsv')
      ruta_comp <- paste0(getwd(), .Platform$file.sep, File_comp, .Platform$file.sep, Name_comp)
      
      if (file.exists(ruta_comp)) {  # Check if file exists
        # Read the file
        my_comp <- read_delim(file = ruta_comp, delim = "\t")
        my_comp <- as.data.frame(my_comp)
        
        # Process data using dplyr
        Result <- my_comp %>%
          group_by(`Row in evidence file`) %>%
          mutate(Min_Distance_Group = min(`Distance (A)`)) %>%
          filter(`Distance (A)` - Min_Distance_Group <= Min_Distance) %>%
          ungroup()
          if (Min_Distance==0){
            Result <- Result %>%
              distinct(`Row in evidence file`, `Distance (A)`, .keep_all = TRUE)
          }
        
        # Write the result to an Excel file
        output_path <- paste0("X:/NICOLAS/Documents/Trabajo/Doctorado-CEA/Bureau_De_Trabajo/R_Coding/R_Analisys/Distance_Clean/", Folder1, '/ShortSubset_',PDB_Model[Z], Treat_Name[X],'.xlsx')
        write_xlsx(Result, output_path)
      } else {
        message(paste("File not found:", ruta_comp))
      }
    }
  }
}
